# O-glycosylation contributes to mammalian glycoRNA biogenesis

**DOI:** 10.1101/2024.08.28.610074

**Authors:** Jennifer Porat, Christopher P. Watkins, Chunsheng Jin, Xixuan Xie, Xiao Tan, Charlotta G. Lebedenko, Helena Hemberger, Woojung Shin, Peiyuan Chai, James J. Collins, Benjamin A. Garcia, Daniel Bojar, Ryan A. Flynn

**Affiliations:** Stem Cell Program and Division of Hematology/Oncology, Boston Children’s Hospital, Boston, USA; Department of Stem Cell and Regenerative Biology, Harvard University, Cambridge, USA; Proteomics Core Facility at Sahlgrenska Academy, University of Gothenburg, Gothenburg, Sweden; State Key Laboratory of Genetic Engineering, Greater Bay Area Institute of Precision Medicine (Guangzhou), School of Life Sciences and Institutes of Biomedical Sciences, Fudan University, Shanghai 200438, China; Department of Biochemistry and Molecular Biophysics, Washington University School of Medicine, St. Louis, MO, USA; Wyss Institute of Biologically Inspired Engineering, Harvard University, Boston, USA; Division of Gastroenterology, Massachusetts General Hospital, 55 Fruit Street, Boston, MA 02114, USA; Harvard Medical School, 25 Shattuck St., Boston, MA 02115, USA; Institute for Medical Engineering & Science and Department of Biological Engineering, Massachusetts Institute of Technology, Cambridge, MA 02139, USA; Department of Bio and Brain Engineering, Korea Advanced Institute of Science and Technology, Daejeon, Republic of Korea; Infectious Disease and Microbiome Program, Broad Institute of MIT and Harvard, Cambridge, MA 02142, USA; Department of Chemistry and Molecular Biology, University of Gothenburg, Gothenburg, Sweden. Wallenberg Centre for Molecular and Translational Medicine, University of Gothenburg, Gothenburg, Sweden; Harvard Stem Cell Institute, Harvard University, Cambridge, USA

## Abstract

There is an increasing appreciation for the role of cell surface glycans in modulating interactions with extracellular ligands and participating in intercellular communication. We recently reported the existence of sialoglycoRNAs, where mammalian small RNAs are covalently linked to N-glycans through the modified base acp^3^U and trafficked to the cell surface. However, little is currently known about the role for O-glycosylation, another major class of carbohydrate polymer modifications. Here, we use parallel genetic, enzymatic, and mass spectrometry approaches to demonstrate that O-linked glycan biosynthesis is responsible for the majority of sialoglycoRNA levels. By examining the O-glycans associated with RNA from cell lines and colon organoids we find known and previously unreported O-linked glycan structures. Further, we find that O-linked glycans released from small RNA from organoids derived from ulcerative colitis patients exhibit higher levels of sialylation than glycans from healthy organoids. Together, our work provides flexible tools to interrogate O-linked glycoRNAs (O-glycoRNA) and suggests that they may be modulated in human disease.

## Introduction

Glycosylation of RNAs is an emerging topic but remains poorly understood in both scope and function. Initially, a metabolic reporter of sialic acid (peracetylated N-azidoacetylmannosamine, Ac_4_ManNAz) was used to discover that small noncoding RNAs from mammalian cells were modified with N-glycans^1^. These N-glycans have a variety of compositions, which include sialic acid and fucose modifications^1,2^. These “sialoglycoRNAs” are presented on the external surface of live cells and can interact with Sialic acid binding Ig-like lectin (Siglec) receptors^1^. Thereafter, sialoglycoRNAs were also found on neutrophils, and treatment of the neutrophils with RNase caused a reduction in the ability for neutrophils to be recruited to sites of inflammation^3^. In addition, RNase treatment of THP-1 monocytes and macrophages impaired their adhesion to epithelial cells, suggesting that cell surface RNAs may function in various cell surface processes^4^. More recently, we reported a covalent linkage between an N-glycan and RNA that occurs via the modified base 3-(3-amino-3-carboxypropyl)uridine (acp^3^U)^5^, providing evidence of a direct RNA-glycan connection.

In the context of the initial glycoRNA discovery, the Ac_4_ManNAz signal was robustly sensitive to PNGaseF cleavage^1^, directing focus towards characterizing N-glycans. Imaging technologies such as sialic acid aptamer and RNA in situ hybridization-mediated proximity ligation assay (ARPLA), which can visualize specific sialoglycoRNAs on the surface of cells, demonstrated loss of signal after PNGaseF cleavage^4^. Using a newly developed glycomics method leveraging Data Independent Acquisition (DIA) mass spectrometry (MS) methods (called glycanDIA) we recently reported the profile of N-glycans released from small RNAs across five primary murine tissues, finding many sialylated and fucosylated species^6^. Consistent with this, work using more traditional Data Dependent Analysis (DDA) MS found similar features of N-glycans released from twelve primary human tissues^2^. Curiously, during the course of our characterization of the acp^3^U-glycan linkage and using RNA optimized periodate oxidation and aldehyde labeling (rPAL), a non-metabolic reporter approach to detect native sialoglycoRNAs, we found that PNGaseF and loss of acp^3^U biosynthesis enzymes had a significant but minor reduction in the total levels of rPAL signal^5^. This suggested that the bulk of the rPAL-labeled sialoglycoRNA signal was not derived from N-glycans.

O-linked glycans are another major type of protein glycosylation. Much of protein O-glycan synthesis is initiated by addition of N-acetylgalactosamine (GalNAc) to serine or threonine residues by one of up to 20 Golgi-localized GalNAc transferases^7^. In contrast to N-glycans, where a single core glycan is linked to the amide nitrogen on asparagine by OligoSaccharyl Transferase (OST), there are 8 major core structures found in mammalian O-glycans. Depending on the available glycosyltransferase genes, the addition of galactose (Gal) to the initial GalNAc (Galβ1→3GalNAc) forms Core-1 structures and are generated (e.g. in HEK293 cells) in the Golgi by C1GALT1 (the core-1 synthase) and its obligate chaperone COSMC (C1GALT1C1)^8^. Core-1 structures (**Figure 1A**) are found across the body while the others are more tissue selective^9,10^. In line with this, previous glycomics of wild type HEK293 cells revealed that the majority of glycans were based on Core-1 and Core-2 structures^11^. The initiating GalNAc or Core-1 structures can be sialylated on the Gal or GalNAc by Golgi-localized sialyltransferases, or N-acetylglucosamine (GlcNAc) can be added to the Core-1 structure by Core-2 synthases to generate a Core-2 O-glycan (**Figure 1A**)^12^. The lack of a common core structure among O-glycans, as well as the lack of an enzyme capable of releasing intact O-glycans, analogous to PNGase F for N-glycan removal (reviewed in ^13^), have made structural studies of O-glycans difficult and to date, no O-glycans have been unambiguously detected on RNA.

**Figure 1.**
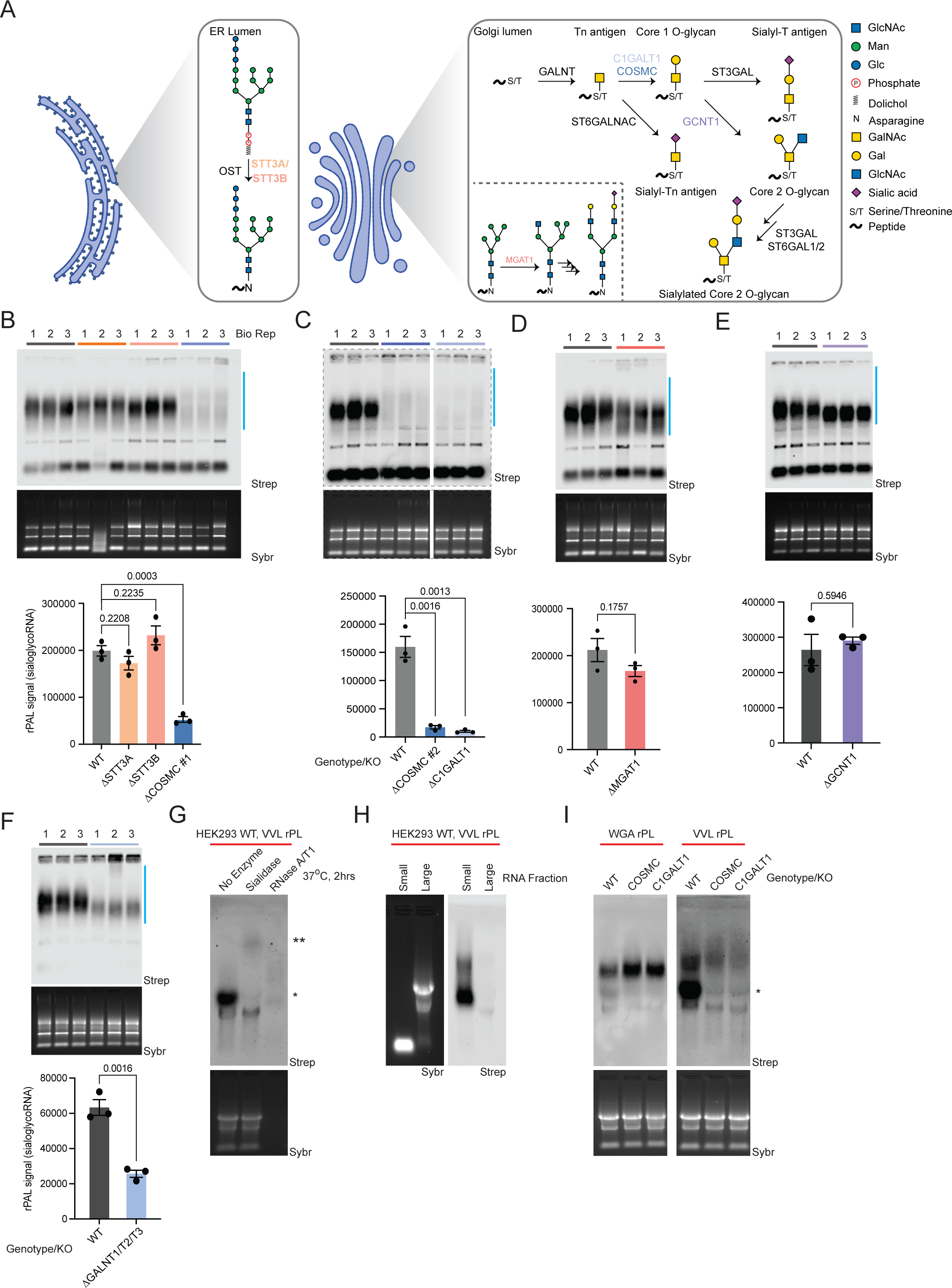
SialoglycoRNAs are dependent O-glycan biosynthetic enzymes. A. Schematic of N-glycan biosynthesis (Endoplasmic Reticulum, Golgi inset) and O-glycan biosynthesis, with the associated glycosyltransferases responsible for each maturation step. B. RNA periodate oxidation and aldehyde ligation (rPAL) northern blot of total RNA from wild type HEK293, STT3A Knockout (KO), STT3B KO, and COSMC KO. An in-gel Sybr stain of total RNA and detection of biotinylated sialoglycoRNA on the membrane (Strep) are shown, along with a quantification of sialoglycoRNA signal (n= 3 biological replicates, mean±SEM, unpaired student’s t-test). C. rPAL northern as in (B), here with WT vs. COSMC KO clone #2 vs. C1GALT1 KO. D. rPAL northern as in (B), here with WT vs. MGAT1 KO. E. rPAL northern as in (B), here with WT vs. GCNT1 KO. F. rPAL northern as in (B), here with WT vs. GALNT1/GALNT2/GALNT3 KO. G. RNA proximity labeling (rPL) of live HEK293 cells using the O-glycan binding lectin VVL (binding GalNAc), followed by *in vitro* digestion of extracted total RNA with sialidase or RNase. *lower molecular weight (MW) signal, ** ultra-high molecular weight signal are noted. H. rPL as in (G), here fractionating small and large RNAs. I. rPL as in (G), here labeling in proximity of VVL (binding a terminal GalNAc) or the N-glycan binding lectin WGA (binding GlcNac and sialic acid) on WT, COSMC KO, or C1GALT1 KO HEK293 cells.

Here we investigate the hypothesis that some of the glycans on RNA may be of an O-linked nature. We use parallel technical approaches to interrogate the composition of O-glycans released from small RNAs isolated from mammalian cells. Using single-gene knockout cell lines and rPAL, we identify the glycosyltransferase genes required for cellular sialoglycoRNA biogenesis. We also present an optimized workflow to label glycoRNAs with galactose oxidase (GAO), expanding the pool of gel-detectable glycoRNAs beyond sialoglycoRNAs. We characterize sialylated and non-sialylated O-glycan structures on mammalian RNA using DIA and DDA mass spectrometry, revealing known and novel O-glycan structures. Finally, we demonstrate that O-linked glycans on RNA from colon organoids derived from patients with Ulcerative Colitis exhibit higher levels of sialic acid than healthy organoids, suggesting that O-linked glycoRNAs may become dysregulated in disease states. Together we uncover O-glycoRNA as a major contributor to the total levels of glycoRNAs in mammalian cells.

## Results

### Genetic dissection of human glycosyltransferases reveals O-glycans as the major component of sialoglycoRNAs

Recent large-scale efforts have been undertaken to examine how each glycosyltransferase (GTase) impacts various aspects of glycan production in human cells^11,14^, however these efforts were focused largely on glycoprotein phenotypes. Here we applied rPAL labeling to 21 individual GTase knockout (KO) HEK293 cells^14^. Importantly, rPAL labels native sialoglycoRNA^5^, thereby eliminating the need for metabolic reporters that may have different sialyltransferase preferences^15^. We initially profiled the rPAL signal generated from total RNA extracted from WT HEK293 cells, OST KOs (STT3A-KO or STT3B-KO, involved in N-glycan biogenesis), or COSMC-KO^8^. Neither STT3A-KO nor STT3B-KO led to a significant loss of overall rPAL signal, while the distribution of rPAL signal from STT3A-KO was slightly up-shifted (**Figure 1B, Table S1**). In contrast, COSMC-KO led to a ∼75% loss in rPAL signal and a broadening of the remaining signal, resulting in a split smear (**Figure 1B**). Loss of COSMC leads to truncated O-glycans and an overall reduction in sialylation^14^, consistent with a loss of sialic acid containing O-glycans on RNA as read out in the rPAL blots. We confirmed here that the loss of C1GALT1 mirrored that of COSMC-KO as well as validated the COSMC phenotype with a second KO clone (**Figure 1C**). Loss of MGAT1, critical for the biogenesis of all known complex N-glycans amenable for sialic acid capping (**Figure 1A**), did not result in a statistically significant reduction in rPAL signal but was accompanied by a minor change in migration in the gel (**Figure 1D, S1A, Table S1**).

Given the apparent contribution of O-glycan biogenesis enzymes to rPAL signal, we examined how Core-1 and Core-2 synthases impacted rPAL signal. HEK293 cells have been shown be dominated by Core-2 structures, with no Core-3 or Core-4 structures on protein templates^11^. COSMC/C1GALT1 are important for generating Core-1 and −2 type O-glycans, so we evaluated the rPAL signal after GCNT1-KO (responsible for Core-2 type O-glycans) and while we found no obvious change in intensity compared to WT, there was a downshifting in the rPAL signal (**Figure 1E, S1B**), suggesting some contribution of Core-2 O-glycans to rPAL signal. Since we saw a loss of rPAL signal in COSMC and C1GALT1 KOs, we would predict that truncated (asialylated) O-glycans remain or are increased in their apparent abundance. Consistent with this, combinatorial KO of COSMC and over expression (OE) of the sialyltransferase ST6GALNAC1 (which can sialylate GalNAc) recovered the rPAL signal lost after KO of COSMC alone (**Figure S1C**). Finally, much of the biosynthetic flux of O-glycosylation occurs through the action of polypeptide N-acetylgalactosaminyltransferase (ppGalNAc-T, GALNT) 1 and 2 with additional contribution from ppGalNAc-T3, which glycosylate serine and threonine on proteins in the secretory pathway^16,17^. Combined KO of ppGALNT1,2,3 resulted in approximately 60% reduction in the rPAL signal (**Figure 1F**), confirming a dependency on cellular, secretory O-glycosylation.

There are 20 known membrane-bound sialyltransferase enzymes that transfer sialic acid to Gal in humans, some of which have specificity for various templates (i.e., lipids vs. proteins^18^). As orthogonal validation of the rPAL signal being due to sialic acid labeling, combined knockout of the sialyltransferases ST3GAL1/2/3/4/5/6 (α2-3 linked sialic acids) and ST6GAL1/2 (α2-6 linked sialic acids) resulted in robust loss of sialoglycoRNA labeling with rPAL (**Figure S1D**). In the background of a complete ST3-KO, knockdown of ST6GAL1/2 did not result in any further changes to rPAL signal (**Figure S1D**). More limited combinations of KO demonstrated partial effects on the rPAL signal, showing that in isolation, loss of ST3GAL4/6, ST3GAL1/2, and ST6GAL1/2 can cause partial reductions in rPAL signal (**Figure S1A, S1B**) and that combining ST3GAL3/4/6 and ST6GAL1/2 KOs can make the rPAL reduction more robust (**Figure S1A**). Consistent with this, single gene KOs of ST3GAL1, 2, or 3 had minimal effect on the total rPAL levels, although knockout of ST3GAL1 or 2 slowed the mobility of sialoglycoRNAs (**Figure S1E**). Taken together, when cells can mostly generate α2-3 linked sialic acids (ST6 loss) the rPAL signal shifts down (faster migration) while when the cells can mostly generate α2-6 linked sialic acid the rPAL signal shifts up (slower migration). Overexpression experiments confirmed this, as over-expression of ST3GAL4 (in a ST3GAL3/4/6 KO) resulted in increased rPAL signal and molecular weight shift down (**Figure S1D**) while over-expression of ST6GAL1 (in a ST3GAL3/4/6, ST6GAL1/2 KO) partially increased the rPAL signal but shifted the molecular weight up (**Figure S1A, Table S1**).

### Small RNAs are in proximity to O-glycans on the cell surface

We previously demonstrated that terminal GlcNAcβ binding lectins like WGA^19^ are in proximity to sialoglycoRNAs on the surface of HeLa cells^1^; however we did not previously assess if O-glycan binding lectins are also in proximity to cell surface sialoglycoRNAs. We next investigated the ability of various biotinylated lectins to label any RNA on the surface of living HEK293 cells (**Methods**). Importantly, these “RNA proximity labeling” (rPL)^20^ experiments monitor the levels of cell surface RNA in proximity to specific lectin ligands, rather than the abundance of the lectin ligand directly. All lectins used were able to label proteins on the surface of WT and KO cells, confirming the binding and labeling activity (**Figure S1F**). Consistent with our previous observations on HeLa cells, the sialic acid-binding lectins WGA and MAA-II generated high molecular weight (MW) biotin signal (**Figure S1G**). We also examined the labeling pattern surrounding ligands of Jacalin (mainly binding Core-1 O-glycans in HEK293 cells^19^) and VVL (GalNAc-binding, Tn antigen^19^) and found that both lectins clearly produced high MW signal, with Jacalin being more similar to WGA and MAA-II. VVL and WGA labeling also produced an ultra-high MW signal (**) as well as signal closer to the 28S region (**Figure S1G**, lower MW, *).

Given that VVL has high specificity for the truncated Tn antigen (terminal GalNAc) O-glycan structure^19^ and displayed a different pattern for high MW biotin signal, we investigated if this biotin signal was related to sialoglycoRNAs. Notably, the rPL assay reports on RNA species in physical proximity to ligands of the utilized lectin (here VVL) on the cell surface. *In vitro* RNase digestion reduced the signal of all bands, including the high MW band and putative rRNA bands (**Figure 1G**). *In vitro* sialidase treatment of the VVL-rPL signal selectively reduced the signal near the 28S rRNA region while maintaining the ultra-high MW region, suggesting that some of the RNA species proximal to VVL substrates are likely sialoglycoRNA (**Figure 1G**). Further, the high MW and 28S-adjacent signal fractionated with the small RNAs and not the long RNAs, consistent with our previous findings that N-glycans are found on small RNA^1^ (**Figure 1H**). We repeated the proximity labeling using WGA and VVL on WT, COSMC-KO, and C1GALT1-KO^14^ HEK293 cells. C1GALT1-KO cells were included to assess if the catalytic activity of O-glycosylation is required or if this observation relates to a non-O-glycosylation activity of COSMC. Proximity labeling around WGA was minimally affected by the loss of either COSMC or C1GALT1 (**Figure 1I**). In contrast, RNA labeled in proximity to VVL was affected by the loss of COSMC and C1GALT1, specifically the disappearance of the lower-MW biotin signal (*) (**Figure 1I**). COSMC or C1GALT1 KO lead to O-glycan truncation and loss of some mature, sialylated O-glycans^8^, which is consistent with the loss of the sialidase-sensitive lower-MW biotin signal (*) in the COSMC and C1GALT1 KO. Truncated mature surface-presented O-glycans in the KO cell lines are still accessible to VVL, leading to the maintenance of higher MW, more poorly sialylated proximal RNA. Together, these data suggest that small RNAs are in close proximity to O-glycans on the cell surface.

### Galactose oxidase labels glycoRNA

Although our findings indicate that the O-glycan biogenesis pathway contributes to sialoglycoRNA levels, many O-glycan structures lack sialic acid in HEK293^11^ (**Figure 1A**), so we sought out an alternative labeling method to expand our detection of glycoRNA beyond sialoglycoRNAs and allow derivatization of another monosaccharide. Recent studies have proposed the use of galactose oxidase (GAO) for labeling glycoRNA^21,22^, although biochemical evidence of GAO’s ability to selectively label glycoRNAs has not been demonstrated yet. GAO is a fungal enzyme that oxidizes galactose (Gal) and *N*-acetylgalactosamine (GalNAc) at the C6 position^23^. Reasoning that this would allow us to label Gal and GalNAc on O-glycans (**Figure 2A**), we optimized a set of conditions that would allow robust labeling while minimizing RNA degradation. Many sources of commercial GAO exist from multiple fungal species and with a wide variety of expression and purification methods; it was therefore necessary to investigate their ability to selectively oxidize glycoRNA *in vitro*. Indeed, when we tested three commercial sources of *Dactylium dendroides* GAO (from Worthington, MedChem Express, and Sigma, whole protein analysis of the commercial stocks in **Figure S2A**), we observed that Worthington GAO specifically generated a product in the presence of small RNA, while MedChem Express GAO generated the same product regardless of whether small RNA was present or absent in the reaction (**Figure 2B**). Although the signal generated by Sigma GAO was RNA-dependent, labeling was substantially weaker than that observed with Worthington, so all subsequent optimizations were performed using Worthington GAO.

**Figure 2.**
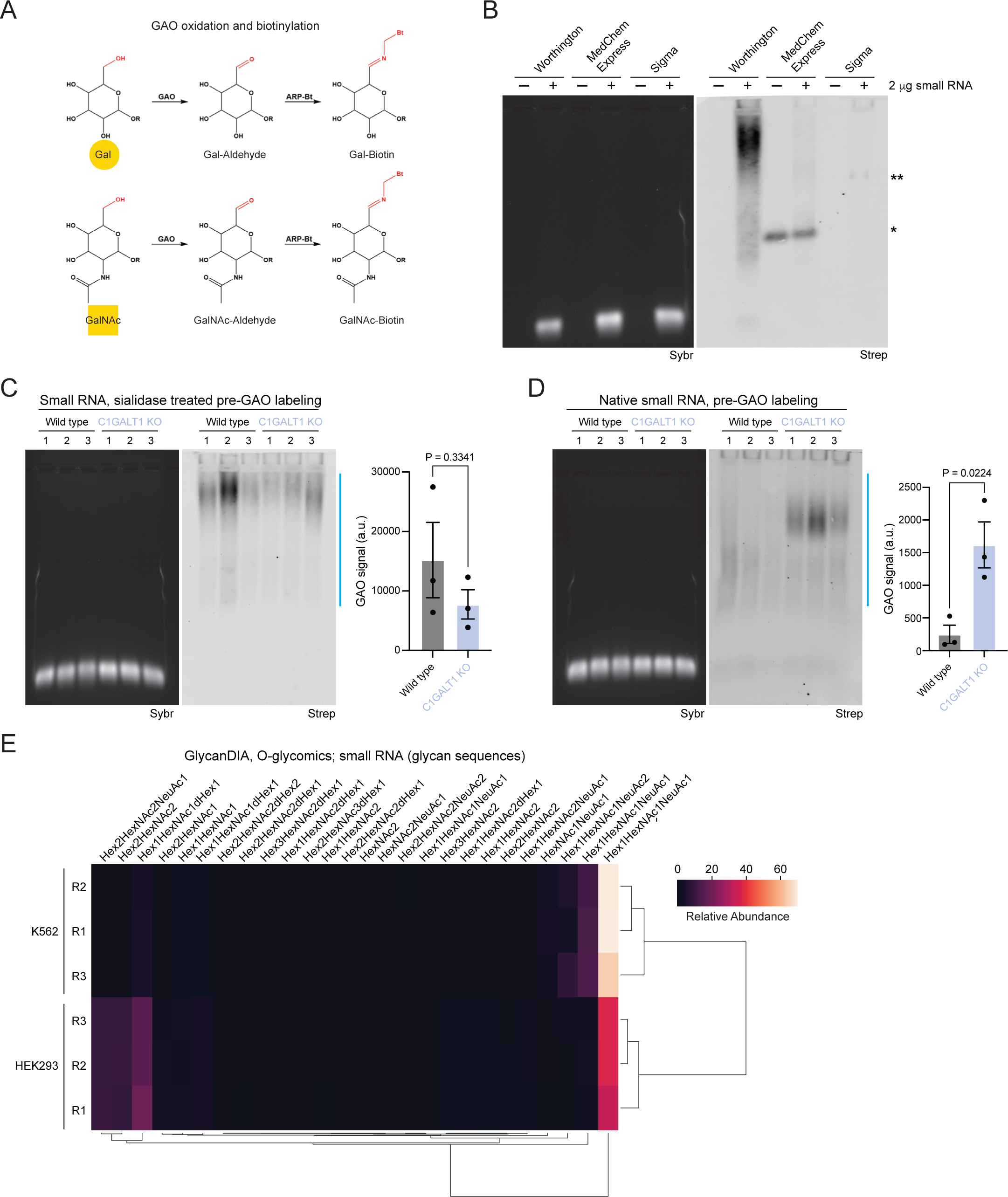
Galactose oxidase (GAO) labels glycoRNA. A. Reaction schematic of the oxidation of galactose (Gal) or *N*-acetylgalactosamine (GalNAc), followed by ligation of biotin (ARP-Bt) to the newly generated aldehyde. B. GAO northern with and without wild type HEK293 small RNA labeled with GAO purchased from Worthington, MedChem Express, or Sigma. *RNA-independent labeling using MedChem Express GAO, **RNA-dependent labeling using Sigma GAO. An in-gel Sybr stain of small RNA and detection of biotinylated glycoRNA on the membrane (Strep) are shown. C. GAO northern as in (B), with wild type and C1GALT1 KO HEK293 small RNA pre-treated with sialidase. A quantification of GAO signal is shown (n= 3 biological replicates, mean±SEM, unpaired student’s t-test). D. GAO northern as in (C), with native (non sialidase-treated) wild type and C1GALT1 KO HEK293 small RNA. E. Heatmap of O-glycan structures released from K562 and HEK293 small RNA and identified by Data Independent acquisition (DIA). Glycan structures are indicated (n= 3 biological replicates).

GAO is a Cu(II) metalloenzyme and a potent oxidizer^23^, to the point that addition of GAO itself is sufficient to degrade small RNA (**Figure S2B**). The addition of RNaseOUT, EDTA, or heat inactivation of GAO failed to rescue RNA degradation, suggesting that degradation is likely not due to contaminating RNase in GAO or GAO catalytic activity (**Figure S2B**). To mitigate the propensity of GAO to degrade small RNA, we titrated the amount of GAO and used a reaction buffer consisting of 20% DMSO and 100 U of HRP per 1 U of GAO (**Figure S2C**), which enhance reaction conditions and rate through multiple mechanisms^24^. We optimized the reactivity of HRP by using a 50 mM phosphate buffer of pH 6, ensuring robust protection of small RNA integrity through high HRP activity (**Figure S2C**). Importantly, Gal and GalNAc are commonly present as terminal and penultimate carbohydrates in N- and O-glycans. When Gal and GalNAc are penultimate carbohydrates, they are linked to the terminal sialic acid at either carbon 3 or 6, with the frequency of which linkage being variable across cell type and cell status. The α2-6 sialic acid linkage directly prevents oxidation by GAO, requiring removal of sialic acid. We therefore introduced a sialidase step prior to the GAO reaction for all samples when establishing our optimized reaction conditions. Finally, *in vitro* RNase digestion prior to GAO labeling resulted in a loss of signal, confirming that our labeling conditions are specific for RNA (**Figure S2D**).

Next, we tested the ability of GAO to detect differences in glycoRNA resulting from O-glycan truncation by comparing GAO labeling of RNA from wild type and C1GALT1 KO cells (**Figure 2C**). We detected no significant difference when we included a sialidase digestion prior to GAO labeling, consistent with the idea that any changes in accessibility of the C6 position on Gal or GalNAc as a result of decreased sialylation in the C1GALT1 KO would be lost upon sialidase treatment of small RNA. In contrast, omitting the sialidase step to enable profiling of native, intact glycoRNA led to increased GAO labeling of the C1GALT1 KO relative to wild type RNA (**Figure 2D**). Taken together, our optimized GAO labeling reaction is suitable for profiling mammalian glycoRNA and yields additional biological insight into the sialylation state of glycoRNA upon comparison of signal with and without prior sialidase treatment.

### O-glycomics reveal an expanded set of glycans that purify with RNA

As an orthogonal means to examine O-glycans on RNA, we purified small RNA from the HEK293 (kidney) and K562 (lymphoblast) immortalized cell lines and used O-glycomics to profile glycans released from small RNA. Because N-glycan profiles have now been established, we focused on possible O-glycan signals. O-glycans were released from small RNA material using standard reductive beta-elimination procedures and processed for MS using positive mode ionization and Data Independent Acquisition (DIA) scans (glycanDIA^6^). In general, the glycoRNAs of HEK293 and K562 cells were well separated and samples clearly clustered by cell line (**Figure 2E, Table S2**), with very few structures shared across all samples, namely sialylated Core-1 O-glycans. We identified a cluster of sialylated Core-2 structures that were characteristic of HEK293 cells, while K562 exclusively presented Core-1 O-glycans on their glycoRNAs, similar to what was seen in a genome-wide knockout screen of Siglec-7 ligands on K562 cells^25^. The finding of Core-1 and −2 structures on HEK293 cells matches those found in protein-linked O-glycans in that cell line^26^. Similarly, for K562 cells, lectin microarray studies^27^ support the prevalence of Core-1 O-glycans, where it should be noted that this measurement likely also inadvertently co-measured glycoRNAs on the surface of these cells.

### O-glycomics on human colon organoids and primary colon tissue RNA

The diversity of N-glycans in glycoRNAs from primary tissues was greater than what we initially observed from cell lines^1,6^, To extend our analysis of glycoRNA beyond 2D cell culture, we set out to profile O-glycans on RNA isolated from 3D colon organoids derived from healthy colon as well as from Ulcerative Colitis (UC) patients^28^, which would also allow us to model a disease process. First, we examined the gel patterns of the sialoglycoRNAs between colon cancer cells (Caco2) and an organoid derived from non-inflamed colon tissue. Here we found a dramatic increase in intensity and a down-shifting of the rPAL signal, per mass of recovered total RNA (**Figure 3A**). The highly abundant and broader smearing rPAL signal observed from the organoid RNA was RNase sensitive (**Figure S3A**), as we would expect for sialoglycoRNAs. We then performed rPAL labeling and O-glycomics on RNA from colon organoids derived from 3 healthy patients (**Figure 3B, C**). Here we used negative mode ionization, given the higher abundance of rPAL signal and thus negatively charged sialoglycans, Data Dependent Acquisition (DDA), and glycosidase treatments to characterize O-glycan structures from organoid small RNA (**Figure 3C**). For DDA glycomics, sialidase treatment of released O-glycans prior to glycomics measurements further enhanced the structural elucidation of the neutral backbones of sialylated glycans, which otherwise only presented uninformative general sialic acid fragmentation products (e.g., B1 at *m/z* 290). Analysis of glycan compositions of O-glycans released from RNA revealed that many compositions have sialic acid (NeuAc) (**Figure 3C**), in agreement with the intense rPAL signal observed for organoid sialoglycoRNA.

**Figure 3.**
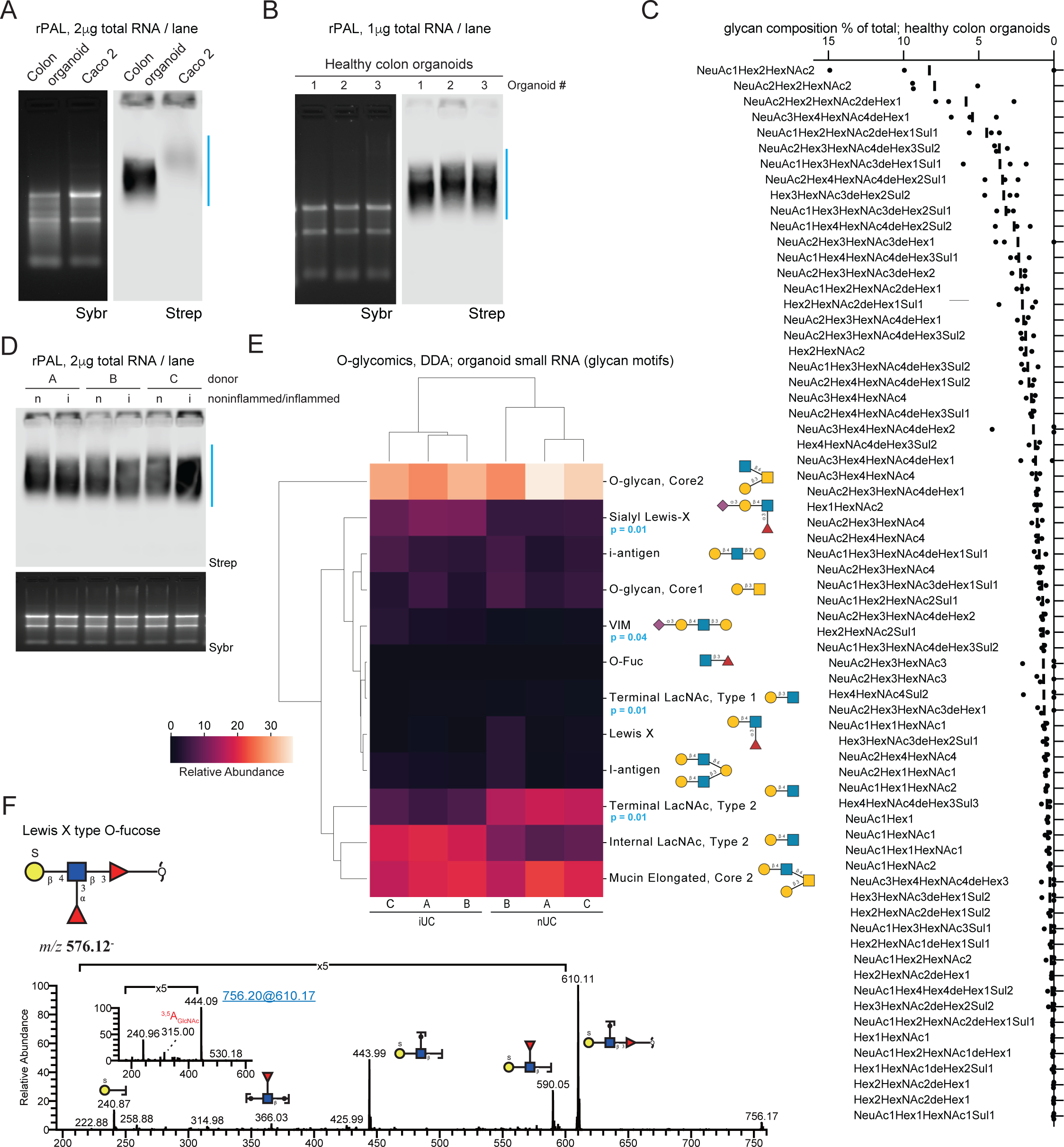
Mass spectrometry defines the O-glycan profile of glycoRNAs. A. rPAL northern blot of total RNA from a colon organoid derived from healthy cells and colon cancer cells (Caco 2). An in-gel Sybr stain of total RNA and detection of biotinylated sialoglycoRNA on the membrane (Strep) are shown. B. rPAL northern as in (A), here with total RNA from colon organoids derived from healthy patients (n= 3 biological replicates). C. Glycan composition percent (for all compositions with a percent greater than 0) of O-glycans released from RNA from healthy organoids and identified by Data-Dependent Acquisition (DDA) (n= 3 biological replicates). D. rPAL northern as in (A), here with total RNA from colon organoids derived from non-inflamed (n) or inflamed (i) cells (n= 3 biological replicates). E. Heatmap of O-glycans identified by DDA in small RNA from organoids derived from inflamed Ulcerative Colitis (iUC) and non-inflamed Ulcerative Colitis (nUC) cells (two-tailed Welch’s t-test corrected by the Benjamini-Hochberg procedure, n= 3 biological replicates). F. MS/MS spectrum of a novel sulfated O-fucose, SGalβ1-4(Fucα1-3)GlcNAcβ1-3Fucα (m/z 576).

To understand whether this pattern changed in the context of disease, separate organoids were generated using cells from inflamed (iUC) and non-inflamed (nUC) regions of the colon from the same patient. rPAL analysis of small RNA from these organoids demonstrated similar total levels but a small down-shifting in the glycoRNA signal in the inflamed samples as compared to the non-inflamed regions (**Figure 3D**). These samples were analyzed for their RNA-linked O-glycans using DDA-MS, detecting 105 unique O-glycans (**Table S3, Figure S3B**); for DDA we were able to structurally characterize 56 glycans (**Table S3**). Consolidating this analysis into glycan motifs^29^, we noticed that nearly all of the glycans were O-GalNAc and O-Fucose (Fuc) type glycans, as well as a few peeling-like structures with hexose at the reducing end (peeling products from beta-elimination reviewed in ^13^), terminating in galactose (**Figure 3E, Table S3**). We next evaluated if there were any statistical differences in the glycan motifs in the inflammatory environment of UC on glycoRNAs, similar to what has been reported for other types of glycans^30^. We performed a differential glycomics expression and identified a shift in the glycome, with a separate clustering of samples from healthy and inflamed tissue (**Figure 3E**, blue stars). This clustering was mainly driven by significantly higher levels of sialylated glycoRNAs in organoids from inflamed tissue (e.g., glycoRNAs exhibiting sialyl-Lewis X and VIM motifs), while the glycoRNAs from normal tissue predominantly exposed undecorated terminal LacNAc units that were sialylated in the inflamed case. This is in agreement with our rPAL results showing a slight downshifting of glycoRNA from inflamed organoids (**Figure 3D**), as sialyl-Lewis X motifs contain α2-3 sialic acid linkages, which migrate faster than α2-6 linkages (**Figure S1B**, ST6 KO). Overall, the identified structures predominantly comprised Core-2 and Core-1 structures, with very few Core-5 structures (**Table S3**). This is notable, as such a profile would instead be expected from salivary mucin glycans^31^, not the colon from which the tissues have been extracted, and which is typically rich in Core-3 and −4 structures on its protein-linked glycans^32^.

Examining the glycoforms in more detail, we noticed multiple structures derived from our glycoRNA samples that have, to our knowledge, never been described before (using a graph-based search in the glycowork-internal glycan database^33^, accounting for linkage ambiguities). Beyond GalNAc, we found examples of extended O-Fucosylated (Fuc) glycans that exhibited sulfated Lewis structures (e.g., SGalβ1-4(Fucα1-3)GlcNAcβ1-3Fucα) (**Figure 3F, Figure S3C, Table S4**), which has not yet been reported. These sulfated structures could be relevant in the context of the reported increase in Siglec binding that is mediated by sulfation^34^, as we have shown that Siglecs bind to glycoRNAs^1^. In total, we identified 6 O-fucosylated O-glycans (m/z 530, 610, 676, 756, 821, 967), of which 2, m/z 530 and 821, are known glycan structures that have been described in human coagulation Factor IX^35,36^ and Notch1^37^ (**Figure S3, Table S4**). The Y ions at m/z 165 in both MS2 and MS3 (inset) suggests the presence of Fuc at the reducing end and the cross-ring cleavage at m/z 263 indicated a β1,4-linked terminal Gal residue (**Figure S3D**). Our observation of a Lewis X type O-fucose is new, where fragmentation ions at m/z 366 and 590 suggest that one Fuc is attached to an internal *N*-acetylglucosamine (GlcNAc) and another Fuc is at the reducing end (**Figure 3F**). The MS3 of fragment ions at m/z 610 confirms that the sulfated Gal is linked to the 4 position of GlcNAc (3.5A cleavage at m/z 315) (**Figure 3F, Table S4**).

## Discussion

Here we have expanded upon our previous observations that cell surface glycoRNAs are modified with N-glycan structures^1,5^ to demonstrate the dependency of glycoRNA levels on O-glycan biosynthesis and presence of O-glycans on RNAs: O-glycoRNA.

We demonstrate the utility of previously developed tools for glycoRNA analysis, such as native sialoglycoRNA labeling with rPAL^5^, lectin proximity labeling^1^, GAO labeling^21,22^, and glycanDIA^6^, and DDA-MS for profiling O-glycans on glycoRNA. In addition to providing information on sialoglycoRNA abundance—where we have shown that the majority of rPAL signal from cancer cell lines comes from O-glycans—rPAL can also provide information on glycan composition through qualitative and quantitative analysis of gel migration. The nature of the sialic acid linkage influences the apparent molecular weight of the glycoRNA signal, with α2-6 sialic acid linkages migrating slower than α2-3 linkages (**Figure 1**). These gel-based insights, coupled with systematic analysis of the effects of knocking out human glycosyltransferases, have thus revealed that sialoglycoRNAs detected by rPAL are chemically diverse and heterogeneous in nature, consisting of N-glycans^1,5^ and O-glycans, which contain α2-3 and α2-6 sialic acid linkages. Importantly, this composition is specific to HEK293 cells, as our O-glycomics revealed unique glycoRNA profiles across cancer cell types and human colon organoids and tissues (**Figure 3**). It will therefore be interesting to assess the relative contribution of different glycosyltransferases to rPAL and GAO signals from different cell lines, cell types, and tissue samples to continue understanding how different glycan biosynthetic pathways result in cell type-specific glycoRNA profiles.

Previous work demonstrated GAO-mediated labeling of total RNA, hydrazide bead coupling, and PNGaseF release to analyze N-glycoRNA^21^ and more recently it was reported that O-glycosidase treatment of the hydrazide beads leads to release of RNA^22^, although the biochemical specificity of GAO in total RNA was not made clear in these works. We note that GAO could also label tRNA containing the hypermodification galactosyl-queosine (GalQ)^38,39^ or other currently known RNA modifications, so specific examination of the labeling patterns of new glycoRNA labeling methods is critical. Here, we have established an updated protocol using GAO to label mammalian glycoRNA and demonstrated its utility for profiling sialylated and asialylated glycoRNA (**Figure 2**). Our method has been optimized for maintaining RNA integrity, which is an important consideration for additional applications, such as sequencing GAO-labeled glycoRNA. Further, we show that truncated O-glycans can be labeled with GAO, and that truncation can be assessed by comparing GAO signal with and without sialidase treatment prior to oxidation. Thus, combined use of rPAL and GAO labeling will enable the characterization of an expanded set of mammalian glycoRNAs. Further, extending these methods for native glycoRNA isolation and coupling this to mass spectrometry could provide evidence of specific linker structures that connect O-glycans directly to mammalian small RNAs.

Lectin blotting is another informative tool for inferring the glycan composition of glycoconjugates including glycoRNA, and repurposing lectins for cell surface proximity labeling provides additional topological insight demonstrating the presence of small RNA proximal to O-glycans on the cell surface. As cell surface RNA has been linked to various immune functions such as neutrophil recruitment to sites of inflammation^3^ and monocyte binding to epithelial cells^4^, future work should address the relative contribution of N-glycosylation and O-glycosylation of RNA using the single gene KOs we have biochemically defined here or inhibitors of various glycosyltransferases.

When comparing the cell line glycans to the colon organoids, we found that the glycans released from the organoids comprise nearly only O-linked glycans (**Figure 3**). It cannot be excluded that small amounts of N-glycans may also be present on these samples, yet it seems clear that these samples have a much greater abundance of O-glycans, analogous to the rich protein-linked O-glycosylation of mucins in this tissue^40^. A possible explanation for this could be a partly shared biosynthetic machinery for glycoproteins and glycoRNA, which predisposes cells in these tissue niches to produce O-glycans on their proteins and RNA. We further note that, even in the case of the cell lines with robustly measurable N-glycans on their RNA, O-glycans still dominated the total signal.

The restructuring of the RNA O-glycome in the comparison of inflamed and uninflamed tissue regions, with statistically significant and interpretable differences even at very small sample sizes, indicates the potential of this new class of biomolecules as a source for diagnostic biomarkers and potential therapeutic intervention. We envision that diagnostic information derived from glycoRNAs could be synergistic in combination with glycomic information from other parts of the glycome, for instance enabling the future development of glycan-based risk scores that integrate glycoproteomics and glycanDIA and DDA for glycoRNA. Given the potentially related aspects of glycan biosynthesis between protein- and RNA-linked glycans, future work should elucidate which features of the respective glycome are regulated “globally” (all biopolymers) and which are subject to carrier-specific regulation (protein vs. RNA). This would both advance the research of these important molecules as well as identify opportunities for specifically modulating aspects of the glycome for therapeutic purposes.

Individual loss of GTase activity can lead to cell compensation which may alter the flux through the glycosylation pathway outside of the specific gene perturbed. For example, loss of early GTase activity like that with OST or GALNTs can change the expression of other GTases and chaperones that depend on N- and O-glycans for proper biological function. Therefore, the effect of an individual GTase may need to be examined in more depth and with other strategies outside simple KO to fully understand their impact on glycoRNA biosynthesis. More generally, our work here does not achieve a direct understanding of the linkage between the O-glycans and small RNA. While this will be important to fully characterize the biosynthesis and atomic nature of O-glycoRNAs, it is likely that significant technology development will be required to enable this discovery.

## Figure Legends

**Figure S1.**
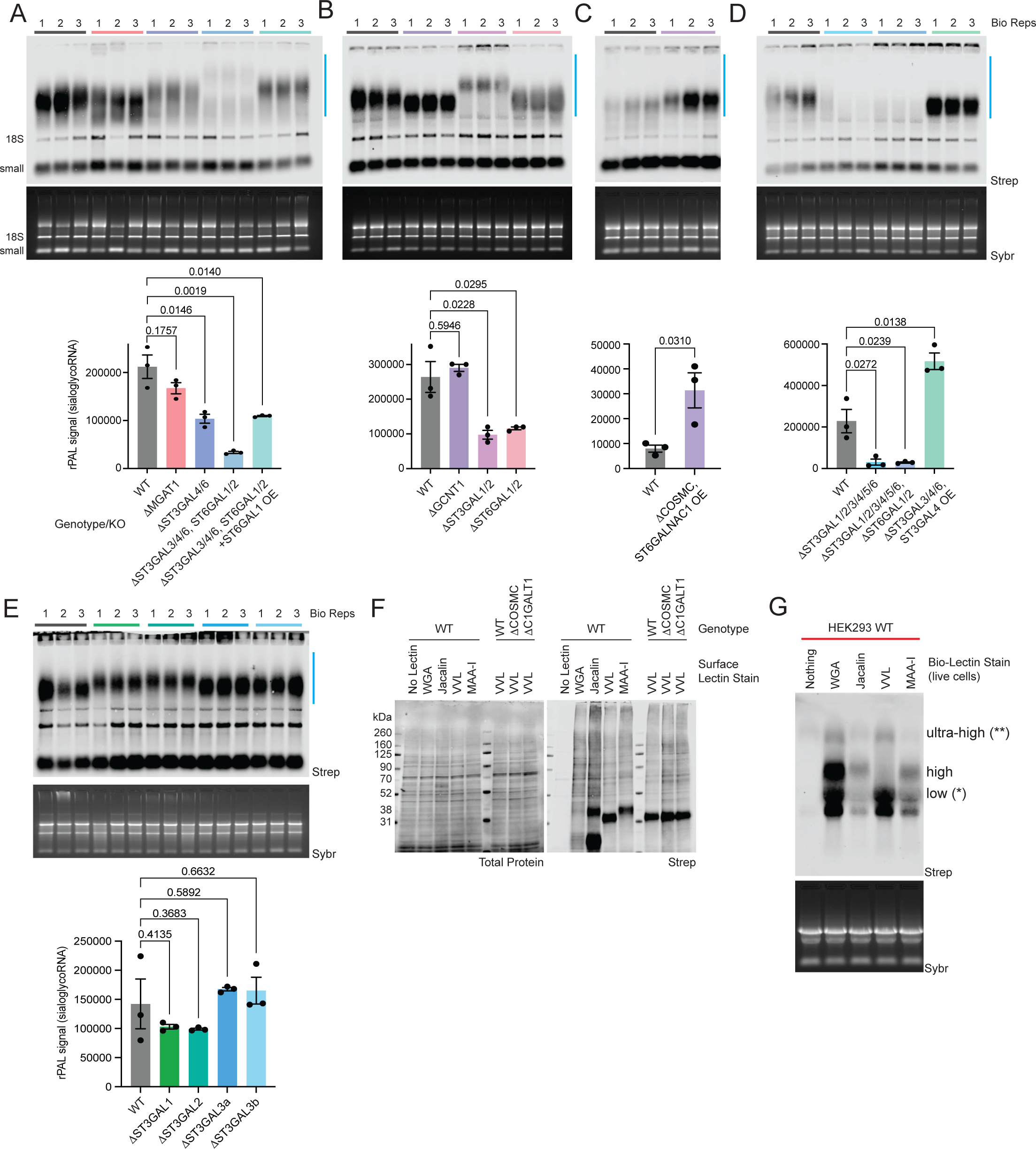
Dissection of O-glycan biosynthesis with sialoglycoRNA labeling. A. rPAL northern with WT, MGAT1 KO, ST3GAL4/6 KO, ST3GAL3/4/6+ST6GAL1/2 KO, and ST3GAL3/4/6+ST6GAL1/2 KO + ST6GAL1 over expression (OE). An in-gel Sybr stain of total RNA and detection of biotinylated sialoglycoRNA on the membrane (Strep) are shown, along with a quantification of sialoglycoRNA signal (n= 3 biological replicates,mean±SEM, unpaired student’s t-test). B. rPAL northern as in (A), here with WT, GCNT1 KO, ST3GAL1/2 KO, and ST6GAL1/2 KO. C. rPAL northern as in (A), here with WT, COSMC KO + ST6GALNAC1 OE. D. rPAL northern as in (A), here with WT, ST3GAL1/2/3/4/5/6, ST3GAL1/2/3/4/5/6+ST6GAL1/2, and ST3GAL3/4/6 KO + ST3GAL4 OE. E. rPAL northern as in (A), here with WT, ST3GAL1 KO, ST3GAL2 KO, ST3GAL3 KO clone a, ST3GAL3 KO clone b. F. Western blot of whole cell lysate from RNA proximity labeling (rPL) experiments. Total protein stains from each sample (left) demonstrate even loading while biotin signal (Strep) demonstrate Lectin-dependent patterns of biotinylation. G. RNA proximity labeling (rPL) northern blot of live HEK293 cells using the following biotinylated lectins: control (no lectin), WGA, Jacalin, VVL, and MAA-I. *lower molecular weight (MW) signal, ** ultra-high molecular weight signal are noted.

**Figure S2:**
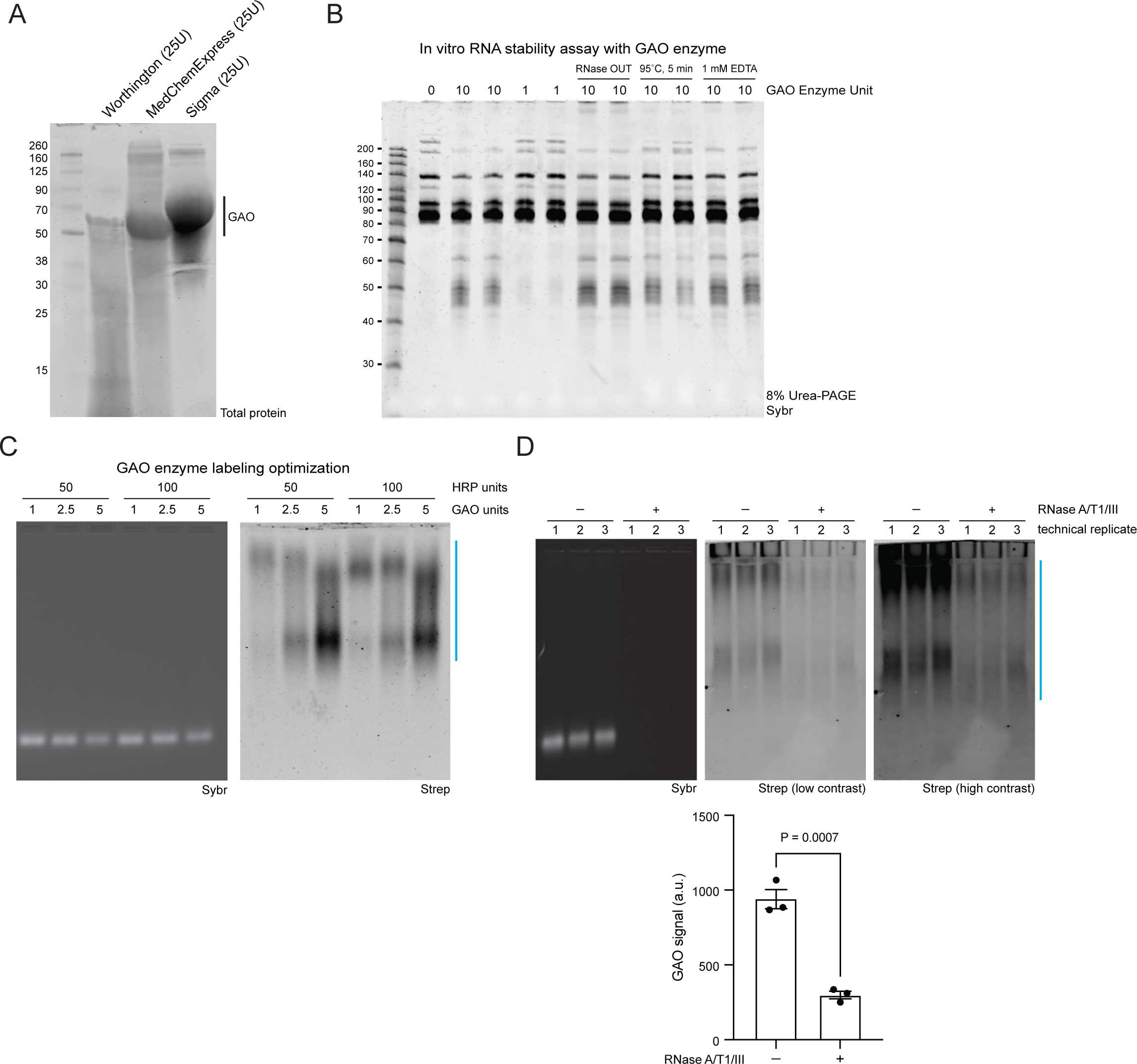
Optimization of GAO for glycoRNA labeling. A. SDS-PAGE of Worthington, MedChemExpress, and Sigma GAO (65-68 kDa). B. Urea PAGE of HEK293 small RNA incubated with 1 or 10 units of Worthington GAO supplemented with RNaseOUT or EDTA, or heat-inactivated GAO. C. GAO northern of HEK293 small RNA treated with various amounts of Worthington GAO and HRP. An in-gel Sybr stain of small RNA and detection of biotinylated glycoRNA on the membrane (Strep) are shown. D. GAO northern as in (B) of GAO-labeled HEK293 small RNA with and without RNase A/T1/III pre-treatment (n= 3 technical replicates, mean±SEM, unpaired student’s t-test).

**Figure S3.**
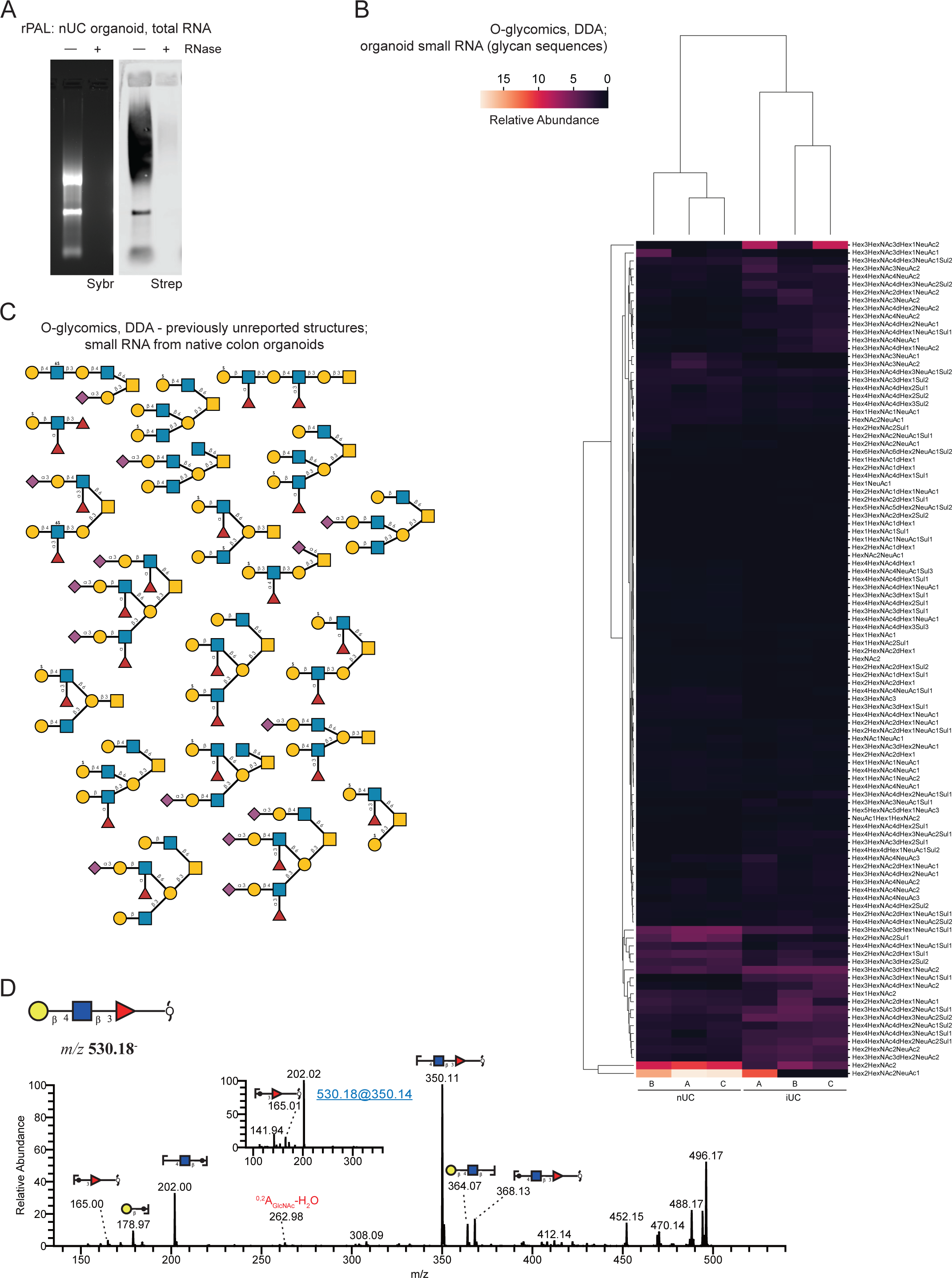
rPAL and mass spectrometry characterization of O-glycans on RNA from Ulcerative Colitis tissue-derived organoids. A. rPAL northern of total RNA from nUC organoids with or without RNase A/T1 treatment. An in-gel Sybr stain of total RNA and detection of biotinylated sialoglycoRNA on the membrane (Strep) are shown. B. Heatmap of O-glycan sequences released from organoids derived from inflamed (iUC) and non-inflamed (nIC) Ulcerative Colitis tissue and identified by Data Dependent Acquisition (DDA) (n=3 biological replicates) C. Previously unreported O-glycan structures that were discovered from the DDA-based MS analysis of O-glycans derived from human colon organoid small RNA. Proposed compositions and linkages are shown and annotated. D. MS/MS spectrum of one conventional O-fucose, Galβ1-4GlcNAcβ1-3Fucα (m/z 530).

**Table S1. Summary of rPAL signal intensity and migration changes upon GTase KOs and OEs.**

**Table S2. Data independent acquisition Mass Spec O-glycomics from HEK293 and K562 cell lines.**

**Table S3. Data dependent acquisition Mass Spec O-glycomics from organoids.**

**Table S4. O-glycan library and GlycanDIA window setup for the analysis.**

## ACKNOWLEDGMENTS

We thank Henrik Clausen, Yoshiki Narimatsu, Sergey Vakhrushev, and Yang Zhang for their generous sharing of clonal knockout cells of the glycosyltransferases as well as extensive and thoughtful discussions of O-glycan biogenesis. We also thank Stacy Malaker for helpful discussions about O-glycan mass spectrometry and other members of the Flynn lab for helpful comments and discussions.

This work was supported by grants from Burroughs Wellcome Fund Career Award for Medical Scientists (R.A.F.), The Rita Allen Foundation (R.A.F.), National Institute of General Medical Sciences of the National Institutes of Health under award number GM151157 (R.A.F.), Branco Weiss Fellowship – Society in Science (D.B.), the Knut and Alice Wallenberg Foundation (D.B.), and the University of Gothenburg, Sweden (D.B.), the American Gastroenterological Association Research Scholar Award (X.T.), NIH NIDDK 1K08DK132516 (X.T.), and the Wyss Institute Clinical Fellowship (X.T.). We thank SciLifeLab and BioMS, funded by the Swedish research council, for providing financial support to the Proteomics Core Facility, Sahlgrenska Academy.

## AUTHOR CONTRIBUTIONS

R.A.F. conceived the project. R.A.F., D.J., J.J.C., and B.A.G. supervised the project and obtained funding. R.A.F., J.P., H.H., C.G.L., and P.C. performed glycoRNA labeling experiments and RNA isolation. C.P.W. designed and optimized the GAO labeling method and performed the glycoRNA labeling experiments with J.P., W.S., and X.T. derived, cultured, and processed colon organoids. Y.X. performed glycanDIA experiments and analyzed the data with B.A.G. J.C. performed DDA glycomics experiments and analyzed the data with D.B. R.A.F. and J.P. wrote the manuscript. All authors discussed the results and revised the manuscript.

## Methods

### Data and code availability

- Raw Mass Spectrometry data have been deposited in the GlycoPOST database and accession numbers are listed below

|  |  |  |
| --- | --- | --- |
| Mass Spectrometry Data for organoids (Data Dependent Acquisition) | This Study | GlycoPOST<br>GPST000463 |
| Mass Spectrometry Data for HEK293 and K562 (Data Independent Acquisition) | This Study | GlycoPOST,<br>GPST000455 |

### Cell culture

HEK293 and K562 (ATCC) cells were grown at 37°C and 5% CO_2_. HEK293 cells were cultured in DMEM media (ThermoFisher Scientific) supplemented with 10% fetal bovine serum (FBS, ThermoFisher Scientific) and 1% penicillin/streptomycin (P/S, ThermoFisher Scientific). K562 cells were cultured in RPMI-1640 media (ThermoFisher Scientific) supplemented with 10% FBS and 1% P/S. All HEK293 KO cells were a gift from the Copenhagen Center for Glycomics^14,41^. All cells were checked for and maintained as mycoplasma free.

### Colon organoid culture

Human colon organoids were grown in 3D culture and harvested as detailed in ^28^. In brief, human colon organoids previously generated from de-identified patient specimens (gift of Dr. Hyun Jung Kim) were removed from liquid nitrogen and thawed at 37°C. 500 µL thawed organoids were resuspended into 9 mL IntestiCult Human Organoid Growth Medium (Stemcell technologies) and centrifuged at 150 g at 4°C for 5 minutes. After removal of supernatant, ice cold Matrigel (Corning) was used to resuspended the organoids (30 µL Matrigel for each 100-200 organoids). The organoids in Matrigel were then plated onto 24-well sterile tissue culture plates (Falcon 353047) on ice with ∼30 µL per well. The plates were inverted to allow 3D Matrigel domes to set for 10 minutes at 37°C. Then, 500 µL of IntestiCult growth medium were added to each well, and organoids were continuously cultured in a humidified cell culture chamber (18% O_2_, 5% CO_2_). Media was changed every other day, and growth was monitored using a phase-contrast microscope.

When significant cellular debris accumulated in the lumen of the organoids (generally 7-10 days), the organoids were split into subcultures by removing media from each well, adding 500 µL ice cold Cell Recovery Solution (Corning), and incubating on ice for 30 minutes to dissolve Matrigel. Wells of the same experimental group were combined into a 15 mL conical tube and centrifuged at 100 g at 4 °C for 5 minutes. Supernatant was removed and 1 mL of room temperature TrypLE Express (GIBCO) was added to each conical tube. The solution was pipetted up and down 5 times before addition of 9 mL room temperature PBS. The conical tubes were then centrifuged at 100 g at 4°C for 5 minutes to pellet the organoids, and supernatant was removed. 30 µL of ice cold Matrigel per each desired new well was added to the pellet on ice with a goal split ratio of 1:3 to 1:5. Organoids were resuspended in Matrigel with gentle pipetting, and 30 uL was pipetted into each well of new 24-well sterile tissue culture plates on ice. The plates were then inverted and to allow 3D Matrigel domes to set for 10 minutes at 37°C. Then, 500 µL of IntestiCult growth medium were added to each well, and organoids were continuously cultured in a humidified cell culture chamber (18% O_2_, 5% CO_2_) as above.

Once sufficient quantities of organoids were obtained by continuous culture, organoids were harvested from Matrigel by incubation in Cell Recovery Solution as described above and pelleted. The pellets were then resuspended in 1 mL RNAzol RT (Molecular Research Center) and vortexed for 1 minute before being flash frozen in liquid nitrogen.

### RNA isolation, Native sialoglycoRNA labeling (rPAL), and blotting

RNA was extracted with RNAzol RT (Molecular Research Center, Inc.) largely following the manufacturer’s protocol and as we have previously reported^42^. For purification of the aqueous phase, it was transferred to clean tubes and 1.1X volumes of isopropanol was added. The RNA is then purified over a Zymo column (Zymo Research). First, 350 μL of pure water was added to each column and spun at 10,000x g for 30 seconds, and the flowthrough was discarded. Next, precipitated RNA from the RNAzol RT extraction (or binding buffer precipitated RNA, below) is added to the columns, spun at 10,000x g for 10-20 seconds, and the flowthrough is discarded. This step is repeated until all the precipitated RNA is passed over the column once. Next, the column is washed three times total: once using 400 μL RNA Prep Buffer (3M GuHCl in 80% EtOH), twice with 400 μL 80% ethanol. The first two spins are at 10,000x g for 20 seconds, the last for 30 sec. The RNA is then treated with Proteinase K (Ambion) on the column. Proteinase K is diluted 1:19 in water and added directly to the column matrix, and then allowed to incubate on the column at 37°C for 45 min. The column top is sealed with either a cap or parafilm to avoid evaporation. After the digestion, the columns are brought to room temperature for 5 min. Next, eluted RNA is spun out into fresh tubes and a second elution with water is performed. To the eluate, 1.5 μg of the mucinase StcE (Sigma-Aldrich) is added for every 50 μL of RNA, and placed at 37°C for 30 minutes to digest. The RNA is then cleaned up again using a Zymo column. Here, 2 volumes of RNA Binding Buffer (Zymo Research) was added and vortexed for 10 seconds, and then 2 volumes (samples + buffer) of 100% ethanol was added and vortexed for 10 sec. This is then bound to the column, cleaned up as described above, and eluted twice with water. The final enzymatically digested RNA is quantified using a Nanodrop under the manufacturer’s RNA settings. For small RNA isolation, 2 volumes of adjusted RNA binding buffer (1:1 Zymo RNA Binding Buffer and 100% ethanol) is added to total RNA and vortexed, followed by binding to a Zymo column. Column flowthrough, consisting of small RNA, is collected, added to 2 volumes of 100% ethanol, vortexed, and bound to a fresh Zymo column.The columns are washed twice with 400 μL 80% ethanol, as above, and eluted twice with water.

rPAL labeling was performed as previously described^42^. Briefly lyophilized, enzymatically treated RNA was blocked. To make the blocking buffer, 1μL 16 mM mPEG3-Ald (BroadPharm), 15 μL 1 M MgSO_4_ and 12 μL 1 M NH_4_OAc pH5 (with HCl) are mixed together (final buffer composition: 570 μM mPEG3-Ald + 500 mM MgSO_4_ + 450 mM NH_4_OAc pH5). 28μL of the blocking buffer is added to the lyophilized RNA, mixed completely by vortexing, and then incubated for 45 minutes at 35°C to block. The samples are briefly allowed to cool to room temperature (2-3 min), then working quickly, 1 μL 30 mM aldehyde reactive probe (ARP/aminooxy biotin, Cayman Chemicals, stock made in water) is added first, then 2μL of 7.5 mM NaIO_4_ is added. The periodate is allowed to perform oxidation for exactly 10 minutes at room temperature in the dark. The periodate is then quenched by adding 3 μL of 22 mM sodium sulfite (stock made in water). The quenching reaction is allowed to proceed for 5 minutes at 25°C. Both the sodium periodate and sodium sulfite stocks were made fresh within 20 minutes of use. Next, the reactions are moved back to the 35°C heat block, and the ligation reaction is allowed to occur for 90 min. The reaction is then cleaned up using a Zymo-I column. 19 μL of water is added to bring the reaction volume to 50 μL, and the Zymo protocol is followed as per the above details. If samples will be analyzed on an agarose gel for glycoRNA visualization, the RNA is then eluted from the column using 2X 6.2μL water (final volume approximately 12 μL).

To visualize the periodate labeled RNA, it is run on a denaturing agarose gel, transferred to a nitrocellulose (NC) membrane, and stained with streptavidin in a manner as per ^42^. After elution from the column as described above, the RNA is combined with 12 μL of Loading Buffer (95% formamide, 18 mM EDTA, 0.025% SDS) with a final concentration of 2X SybrGold (ThermoFisher Scientific) and denatured at 55°C for 10 minutes. Immediately after this incubation, the RNA is placed on ice for at least 2 minutes. The samples are then loaded into a 1% agarose, 0.75% formaldehyde, 1.5x MOPS Buffer (Lonza) denaturing gel. RNA is electrophoresed in 1x MOPS at 115V for between 34 or 45 min, depending on the length of the gel. Subsequently, the RNA is visualized on a UV gel imager, and excess gel is cut away; leaving ∼0.75 cm of gel around the outer edges of sample lanes will improve transfer accuracy. The RNA is transferred as previously described with a buffer of 3M NaCl solution at pH 1, for 90 minutes at 25 °C^42^. Post transfer, the membrane is rinsed in 1x PBS and dried on Whatman Paper (GE Healthcare). Dried membranes are rehydrated in Intercept Protein-Free Blocking Buffer, TBS (Li-Cor Biosciences), for 30 minutes at 25 °C. After the blocking, the membranes are stained using Streptavidin-IR800 (Li-Cor Biosciences) diluted 1:5,000 in Intercept blocking buffer for 30 minutes at 25 °C. Excess Streptavidin-IR800 was washed from the membranes using three washes with 0.1% Tween-20 (Sigma) in 1x PBS for 3 minutes each at 25 °C. The membranes were then briefly rinsed with PBS to remove the Tween-20 before scanning. Membranes were scanned on a Li-Cor Odyssey CLx scanner (Li-Cor Biosciences).

### Galactose oxidase (GAO) labeling and blotting

For the initial sialidase treatment, a 10 uL reaction is performed using 2.5 U of α2-3,6,8 neuraminidase (NEB; Cat. No. P0720L) and 1 U of α2-3,6,8,9 neuraminidase (NEB; Cat. No. P0722L), 50 mM phosphate buffer, pH 6.0, and small RNA (up to 10 ug) for 1 hour at 37 °C. During the final 10 minutes of incubation, both GAO (Worthington; Cat. No. LS004524) and HRP (Sigma-Aldrich; Cat. No. 516531) are reconstituted in UltraPure water (1 U/uL for GAO and 20 U/uL for HRP). Importantly, these reagents must be freshly reconstituted as they are both transition metal-dependent oxidases and lose activity during storage after reconstitution. Following this, 10 uL of the GAO label mix is added, consisting of 100 U of HRP, 1 U of GAO, and 4 uL of DMSO (final concentrations: 5 U/uL HRP, 0.05 U GAO/uL, 20% DMSO). This reaction proceeds at room temperature in the dark for 15 minutes, after which 1 uL of 30 mM ARP-Biotin (Cayman chemicals, Cat. No. 10009350) is added and the reaction incubated at 35 °C for 30 minutes in the dark. Samples were then cleaned using Zymo I columns and eluted using two subsequent volumes of 6.7 uL. For detection by Northern blot, GAO-labeled RNA is loaded into a 1% agarose, 0.75% formaldehyde, 0.8X MOPS Buffer (Lonza) denaturing gel and electrophoresed, transferred, and probed as above.

### In vitro enzyme digestions

All digestions were performed on 30 μg of total RNA in a 20 μL at 37 °C for 60 min. To digest RNA the following was used: 3 μL of RNase cocktail (0.5U/mL RNaseA and 20U/mL RNase T1, Thermo Fisher Scientific) with 20 mM Tris-HCl (pH 8.0), 100 mM KCl and 0.1 mM MgCl_2_. To digest sialic acid: 2 μL of α2-3,6,8 Neuraminidase (50U/mL, New England Biolabs, NEB) with GlycoBuffer 1 (NEB) was added.

### RNA proximity labeling (rPL)

This was performed largely as described previously^1^ with specific modifications noted here. HEK293 cells were cultured as above typically in 10 cm plates. Cells were rinsed once in ice-cold 1x PBS, discarding after each wash, and blocked in Lectin Blocking Buffer (LBB, 20 mM HEPES, 150 mM NaCl, 1 mM MgCl_2_, 1 mM MnCl_2_, 1 mM CaCl_2_, 0.5% BSA) for 15 min at 4 °C. Blocking buffer was then discarded and replaced with LBB+Lectin+Strep-HRP. Typically, 4 mL of this was prepared at a concentration of 5 μg/mL biotinylated lectin and 6 μg/mL Strep-HRP, these components were first mixed together on ice for 30 min prior to the addition of LBB. LBB+Lectin+Strep-HRP staining occurred for 45 min at 4 °C, after which the cells were rinsed twice in ice-cold 1x PBS + 1 mM CaCl_2_ + 1 mM MgCl_2_ (PBS++). Immediately after this 3 mL of PBS++ with 200 μM biotin-aniline (Iris Biotech GMBH) was added to each plate and incubated on ice for 1 min. Plates were then moved to the bench top and H_2_O_2_ was added to a final concentration of 1 mM. This reaction occurred for precisely 2 min, after which plates were brought back on ice, PBS++/ biotin-aniline/ H_2_O_2_ was aspirated, and cells were quickly but gently rinsed twice in Quenching Buffer: 5 mM Trolox, 10 mM sodium ascorbate and 10 mM sodium azide in PBS++. After removing the Quenching Buffer, TRIzol was added directly to the plate and RNA extracted and processed as described above for enzymatic digestions as well as blotting.

### DDA glycomics and analysis

For O-glycan analysis, lyophilized RNA samples were incubated with 50 µL 0.5 M NaBH_4_ and 50 mM NaOH at 50 °C overnight. The released O-glycans were desalted using Dowex 50W. After dryness, the sample was dissolved in 15 µL of water for liquid chromatography-electrospray ionization tandem mass spectrometry (LC-ESI/MS).

The oligosaccharides were separated on a column (10 cm × 250 µm) packed in-house with 5 µm porous graphite particles (Hypercarb, Thermo-Hypersil, Runcorn, UK). The oligosaccharides were injected on to the column and eluted with an acetonitrile gradient (Buffer A, 10 mM ammonium bicarbonate; Buffer B, 10 mM ammonium bicarbonate in 80% acetonitrile). The gradient (0-45% Buffer B) was eluted for 46 min, followed by a wash step with 100% Buffer B, and equilibrated with Buffer A in next 24 min. A 40 cm × 50 µm i.d. fused silica capillary was used as a transfer line to the ion source.

The samples were analyzed in negative ion mode on a LTQ linear ion trap mass spectrometer (Thermo Electron, San José, CA), with an IonMax standard ESI source equipped with a stainless steel needle kept at –3.5 kV. Compressed air was used as nebulizer gas. The heated capillary was kept at 270°C, and the capillary voltage was –50 kV. Full scan (m/z 380-2000, two microscan, maximum 100 ms, target value of 30,000) was performed, followed by data-dependent MS^2^ scans (two microscans, maximum 100 ms, target value of 10,000) with normalized collision energy of 35%, isolation window of 2.5 units, activation q=0.25 and activation time 30 ms). The threshold for MS^2^ was set to 300 counts. Data acquisition and processing were conducted with Xcalibur software (Version 2.0.7).

### DIA glycomics (glycanDIA)

The extracted RNAs were resuspended with 200 µL of 100 mM HEPES buffer, and the mixture was heated using a thermomixer at 100 °C for 2 min. The removal of N-glycans was performed by adding 4 µL of PNGase F (500,000 units/mL, NEB), followed by incubation at 37 °C overnight. The O-glycosylated RNAs were purified by the RNA column (Zymo Research), as noted above. The O-glycans were released using beta-elimination as described in the established protocol^43^. Briefly, the eluted RNAs (45μL) were mixed with 5μL of 2M NaOH and 50μL of 2M NaBH4, and the mixture was incubated at 45 °C for 18 hrs. The reaction was quenched by acetic acid on ice, followed by desalting the O-glycans using the Hypercarb SPE 96-well plate (Thermo Scientific) with 0.1% (v/v) TFA in water washing step and elution with 40% (v/v) ACN and 0.05% (v/v) TFA in water. The eluted O-glycans were dried using the SpeedVac system (Thermo Scientific) and reconstituted with 40% (v/v) ACN and 1% (v/v) TFA in water for further iSPE-HILIC (HILICON) purification. The purified sample was dried using the SpeedVac system and reconstituted in water for LC-MS/MS analysis.

The GlycanDIA analysis was employed as previously described with some modifications for O-glycan analysis^6^. O-glycans were directly characterized using a Waters nanoAcquity UPLC System coupled with a SCIEX ZenoTOF 7600 mass spectrometer. Analytes were separated using Hypercarb Porous Graphitic Carbon HPLC Column (1 mm × 100 mm, 3 μm, Thermo Scientific) at a constant flow rate of 40 μL/min with MS buffer A (0.1% formic acid (v/v) and buffer B (ACN with 0.1% FA (v/v). The chromatography gradient consisted of 0−2 min, 0% B; 2−20 min, 0-16% B; 20−40 min, 16%-72% B; 40−42 min, 72%-100% B; 42-52 min, 100% B; 52-54 min 100%-0% B; 54-70 min 0% B. Ion source gas1, gas2, and curtain gas were at 40, 60, and 35 psi, respectively. O-Glycans were detected under SWATH positive mode, MS1 spectra were acquired over a mass range of *m/z* 300–1650 with 0.25 s accumulation time. The 40 SWATH mass windows were built based on the O-glycan library (**Table S4**), and collision-induced fragmentation was performed with nitrogen gas using dynamic collision energy. The product ions were monitored from the range *m/z* 120-1600 with 20 ms accumulation time.

### Glycomics analysis

The DIA data was analyzed using GlycanDIA Finder^6^ and validated manually using Skyline^44^. All glycans with structural annotations were used to renormalize the relative abundance of each sample to a total sum of 100%. Missing data were imputed from a left-shifted Gaussian distribution (mean = 0.01, standard deviation = 0.01) if group means were below three, otherwise via a k-nearest neighbor imputer, with k = 3. Then, the quantify_motifs function from glycowork (version 0.8) was used with the keyword known, to convert relative abundances of sequences into those of motifs, followed by a renormalization. Variance-based filtering was used to remove motifs with a total variance lower than 1%, followed by variance-stabilizing normalization (i.e., log-transformation and scaling of samples to zero mean, unit variance). Differential expression of motifs then was tested via Welch’s t-tests, followed by a Benjamini-Hochberg correction for multiple testing. All the above is contained within the get_differential_expression function (glycowork version 0.8).

#### Heatmaps for cell line data

Using the relative abundances of O-linked glycans from triplicates of HEK293 and K562 cells, we used the get_heatmap and annotate_figure functions from glycowork (version 0.8.1^33^) to depict the hierarchical clustering of glycan structures rendered in the SNFG style.

#### Heatmaps for organoid data

Heatmaps for organoid glycoRNAs from normal and inflamed tissue from ulcerative colitis patients were generated similar to the ones for cell lines, with the difference of setting “motifs = True” in get_heatmap, choosing the motif subset of “known” within glycowork (version 0.8.1). Differential expression was performed using the get_differential_expression function within glycowork, using the default settings as described in^29^, with the same motif choice as the heatmap. Significantly differentially expressed glycan motifs between organoid glycoRNA from normal and inflamed tissue are depicted on the heatmap, along with their adjusted p-values.

