## Supplementary material for "O-glycosylation contributes to mammalian glycoRNA biogenesis": Table S1

| **Knockout** | **Function** | **Relative signal intensity** | **Relative migration** |
| --- | --- | --- | --- |
| COSMC | O-glycan chaperone for Gal addition | Decreased | Broadening of signal (up-shift and down-shift) |
| C1GALT1 | O-glycan Gal transferase | Decreased | Broadening of signal (up-shift and down-shift) |
| GCNT1 | O-glycan GlcNAc transferase | Unchanged | Down-shift |
| STT3A | Oligosaccharyl transferase (OST) subunit for N-glycan biosynthesis | Unchanged | Up-shift |
| STT3B | OST subunit for N-glycan biosynthesis | Unchanged | Up-shift |
| MGAT1 | Complex N-glycan biosynthesis | Unchanged | Down-shift |
| GALNT1/T2/T3 | O-glycan GalNAc transferase | Decreased | Down-shift |
| ST3GAL4/6 | O-glycan sialyltransferase (α2-3 linked sialic acids) | Decreased | Up-shift |
| ST3GAL3/4/6, ST6GAL1/2 | O-glycan sialyltransferase | Decreased | Broadening of signal (up-shift and down-shift) |
| ST3GAL1/2 | O-glycan sialyltransferase (α2-3 linked sialic acids) | Decreased | Up-shift |
| ST6GAL1/2 | O-glycan sialyltransferase (α2-6 linked sialic acids) | Decreased | Unchanged |
| ST3GAL1/2/3/4/5/6 | O-glycan sialyltransferase (α2-3 linked sialic acids) | Decreased | Down-shift |
| ST3GAL1/2/3/4/5/6, ST6GAL1/2 | O-glycan sialyltransferase | Decreased | Down-shift |
| ST3GAL1 | O-glycan sialyltransferase (α2-3 linked sialic acids) | Unchanged | Up-shift |
| ST3GAL2 | O-glycan sialyltransferase (α2-3 linked sialic acids) | Unchanged | Up-shift |
| ST3GAL3a | O-glycan sialyltransferase (α2-3 linked sialic acids) | Unchanged | Unchanged |
| ST3GAL3b | O-glycan sialyltransferase (α2-3 linked sialic acids) | Unchanged | Unchanged |

| **Knockout** | **Knock in** | **Relative signal intensity** | **Relative migration** |
| --- | --- | --- | --- |
| ST3GAL3/4/6, ST6GAL1/2 | ST6GAL1 | Decreased | Up-shift |
| COSMC | ST6GALNAC1 | Increased | Up-shift |
| ST3GAL3/4/6 | ST3GAL4 | Increased | Down-shift |
